# Predicting chromatin interactions between open chromatin regions from DNA sequences

**DOI:** 10.1101/720748

**Authors:** Fan Cao, Ying Zhang, Yan Ping Loh, Yichao Cai, Melissa J. Fullwood

## Abstract

Chromatin interactions play important roles in regulating gene expression. However, the availability of genome-wide chromatin interaction data is very limited. Various computational methods have been developed to predict chromatin interactions. Most of these methods rely on large collections of ChIP-Seq/RNA-Seq/DNase-Seq datasets and predict only enhancer-promoter interactions. Some of the ‘state-of-the-art’ methods have poor experimental designs, leading to over-exaggerated performances and misleading conclusions. Here we developed a computational method, Chromatin Interaction Neural Network (CHINN), to predict chromatin interactions between open chromatin regions by using only DNA sequences of the interacting open chromatin regions. CHINN is able to predict CTCF- and RNA polymerase II-associated chromatin interactions between open chromatin regions. CHINN also shows good across-sample performances and captures various sequence features that are predictive of chromatin interactions. We applied CHINN to 84 chronic lymphocytic leukemia (CLL) samples and detected systematic differences in the chromatin interactome between IGVH-mutated and IGVH-unmutated CLL samples.

## Introduction

Chromatin interactions play important roles in regulating gene expression^1, 2^. They bridge enhancers to genes^3-5^ and create insulated domains to constrain the reach of enhancers^6^. High-throughput experimental techniques such as high-throughput Chromosome Conformation Capture (Hi-C)^7^ and Chromatin Interaction Analysis with Paired-End Tags (ChIA-PET)^8^ have been developed to detect genome-wide chromatin interactions. These techniques greatly advanced the understanding of genome organization and its roles in transcription regulation^3, 9-11^. However, due to costs and technical challenges, these methods have not been widely applied to large cohorts of cell lines or clinical samples. ChIA-PET requires large amounts of starting materials, limiting its ability to be applied to clinical samples where materials are limited. Hi-C requires extensive sequencing depth to achieve kilobase resolution^12^.

Various computational methods have been developed to predict chromatin interactions to complement the experimental techniques^13-19^. Many of these methods rely on using various functional genomics data including chromatin immunoprecipitation sequencing (ChIP-seq) data of transcription factors and histone modifications, open chromatin data, and transcription data^13, 15, 17, 19^. Methods such as RIPPLE^15^, TargetFinder^17^, and JEME^13^ reported high performances in predicting enhancer-promoter interactions using supervised machine learning approaches. Although the reported performances were exaggerated by using cross-validation with random splitting of samples^20^, these methods suggested that chromatin interactions could be potentially predicted from 1-dimensional functional genomics data^21^. Sophisticated computational methods such as DeepSea^22^ and DeepBind^23^ have demonstrated that many transcription factor binding sites in open chromatin regions could be predicted from DNA sequences. Taken together, these approaches raise the possibility that chromatin interactions between open chromatin regions could be predicted from DNA sequences. A predictor that uses DNA sequences to predict chromatin interactions could potentially expand the understanding of genome organization in new cell types with minimum experimental data and predict the effect of perturbations of non-coding regulatory regions using genome editing techniques such as CRISPR.

In this study, we investigated the possibility of utilizing DNA sequence features to predict chromatin interactions between open chromatin regions. We demonstrated that open chromatin interactions can be predicted accurately from functional genomic data at the resolutions of the experimental techniques. We then developed a novel method to predict open chromatin interactions from DNA sequences and explored the sequence features that were important in predicting chromatin interactions. The models were then applied to a cohort of 84 chronic lymphocytic leukemia (CLL) samples^24^ to investigate the differences between IGHV-mutated and IGHV-unmutated subtypes of the malignancy.

## Results

### Open chromatin interactions can be predicted from functional genomic features

In light of Xi et al. ^20^ and our previous study^21^ showing that the existing prediction methods have exaggerated performances, we first tried to demonstrate that chromatin interactions could be predicted from functional genomic data. Many previous studies focused on enhancer-promoter interactions that were annotated using chromatin interactions derived from Hi-C or ChIA-PET ^13, 15, 17^. The enhancers used were typically hundreds of base pairs, while the chromatin interaction anchors were much larger in size. The resolution discrepancy could lead to the introduction of a lot of noises to the training datasets (Figure 1a). Thus, we used the chromatin interaction anchors directly.

**Figure 1:**
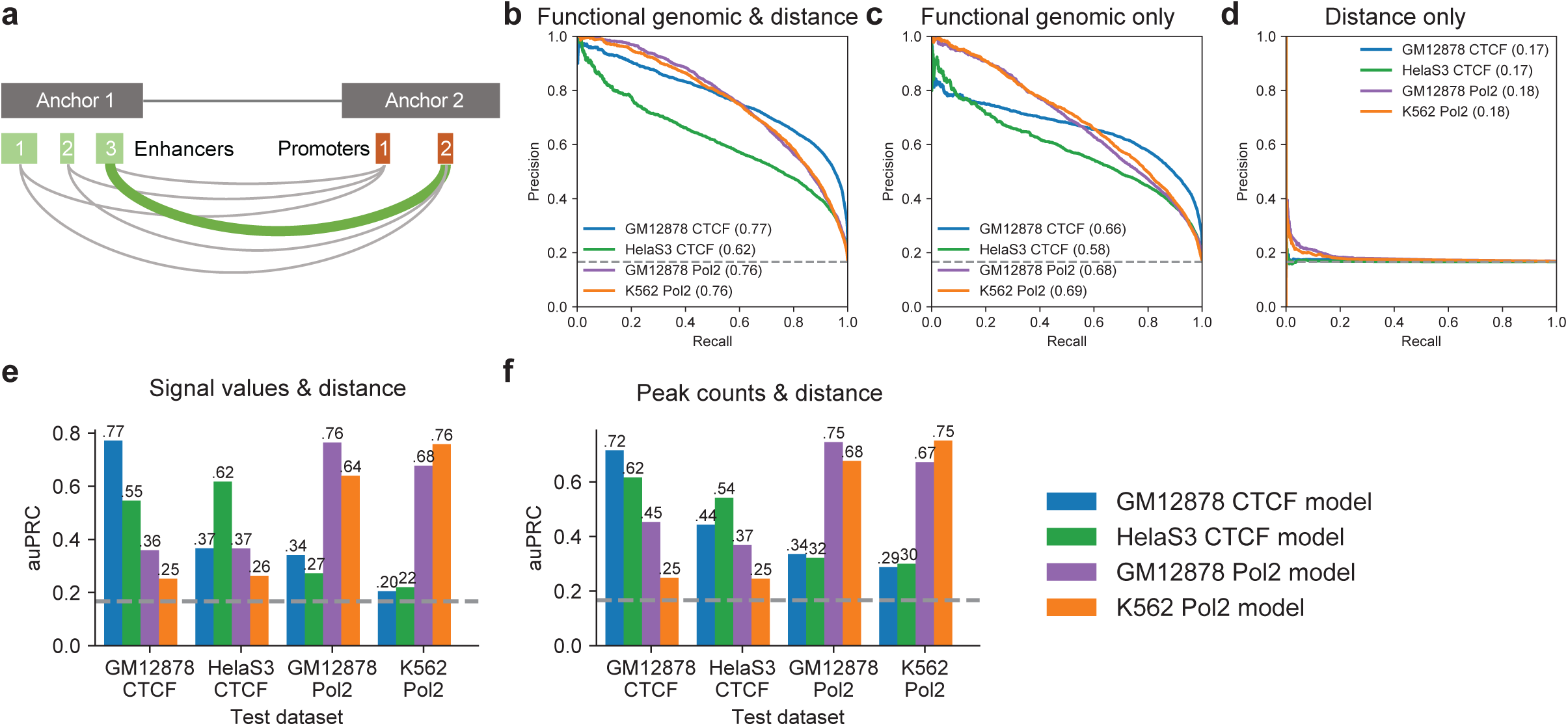
Performances of the functional genomic models on distance-matched datasets. **a**, Illustration of resolution discrepancy between cis-regulatory elements and chromatin interaction anchors. **b-d**, Precision-recall curves of the functional genomic models on distance-matched datasets using features based on b, functional genomic data and distance; **c**, only functional genomic data; and **d**, only distance. Numbers in brackets indicate the area-under precision-recall curve. **e-f**, Across-sample performances using distance and **e**, signal values and **f**, peak counts.

Positive samples were constructed from ChIA-PET datasets separately and the corresponding distance-matched negative datasets were generated. The resulting distance-matched datasets have positive-to-negative ratios of approximately 1:5 and all chromatin interactions were between open chromatin regions in the corresponding cell types. We used ChIP-seq data of transcription factors and histone modifications commonly available to GM12878, K562 and HelaS3 and DNase-seq data from ENCODE^25^ to annotate the anchors and build the feature vectors (Supplementary Table 1). For each chromatin interaction, the average signal of each transcription factor, histone modification and open chromatin were calculated for both anchors. The distance between two anchors was also used as a feature.

Gradient boosted trees^26^ were used to build models for each dataset. We tested three feature sets: 1) all common functional genomics data and distance; 2) distance only; and 3) common functional genomics data only. The models trained on all features achieved area under precision-recall curve (auPRC) ranging from 0.62 to 0.77 (Figure 1b), while models trained on distance are mostly at baseline (Figure 1d), showing that distance is properly controlled between positive and negative samples. The models trained on functional genomics features achieved auPRCs ranging from 0.58 to 0.69 (Figure 1c), lower than models trained on all features. These results showed that although distance alone cannot predict chromatin interactions, the interaction between distance and other features could help to distinguish between positive and negative chromatin interactions.

The across sample performances were lower than within-sample performances (Figure 1e). Using peak counts instead of signal values produced better across-sample performances but lower within-sample performances (Figures 2f). Models trained on RNA Polymerase II (Pol2) datasets generalize well to each other. Models trained on CTCF ChIA-PET datasets, however, did not generalize well to each other. Models trained on CTCF ChIA-PET data perform poorly on Pol2 ChIA-PET datasets and vice versa.

**Figure 2:**
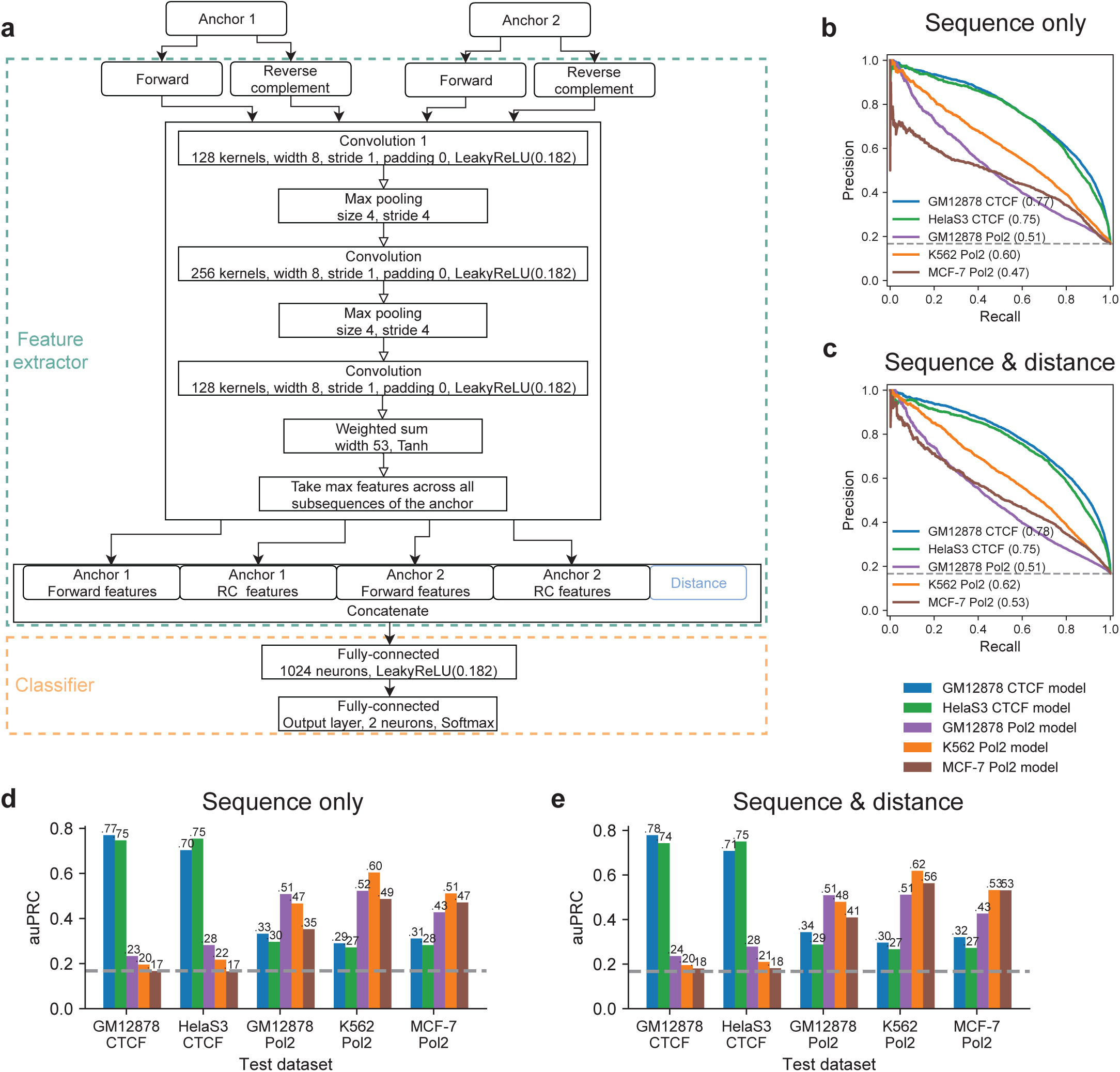
Architecture and performances of the sequence-based models on distance-matched datasets. **a**, The architecture of the sequence-based models using to train on distance-matched datasets. **b-c**, Precision-recall curves of the sequence-based models on distance-matched datasets using b, only sequence features or **c**, sequence features with distance. The numbers in the brackets indicates the area under precision-recall curves. **d-e**, Across-sample performances as measured by area-under precision-recall curve (auPRC) of the models on distance-matched datasets using **d**, only sequence features or **e**, sequence features with distance.

### Open chromatin interactions can be predicted from DNA sequences

Motivated by the results above, we went on to explore whether open chromatin interactions can be predicted from DNA sequences. We built a convolutional neural network, CHINN, to predict chromatin interactions between open chromatin regions using DNA sequences (Figure 2a). The models were trained on GM12878 CTCF, GM12878 Pol2, HelaS3 CTCF, K562 Pol2, and MCF-7 Pol2 datasets separately.

Compared to using functional genomics data for prediction, using sequences produced better within-sample performances for CTCF ChIA-PET datasets with auPRCs of 0.77 for GM12878 CTCF and 0.75 for HelaS3 CTCF (Figure 2b), but worse within-sample performances for Pol2 ChIA-PET datasets with auPRC of 0.51 for GM12878 Pol2, 0.6 for K562 Pol2, and 0.47 for MCF-7 Pol2. Including distance as a feature to classifier only slightly improved the performances for the distance-matched datasets (Figure 2c). The two CTCF models showed good performances on CTCF datasets from both cell types (Figure 2d). Pol2 models also generalize well to other Pol2 datasets. However, models trained on CTCF ChIA-PET datasets perform poorly on Pol2 ChIA-PET datasets and vice versa (Figure 4d-e). The inability to generalize between CTCF models and Pol2 models could be attributed to the different sequence contexts the models captured.

**Figure 3:**
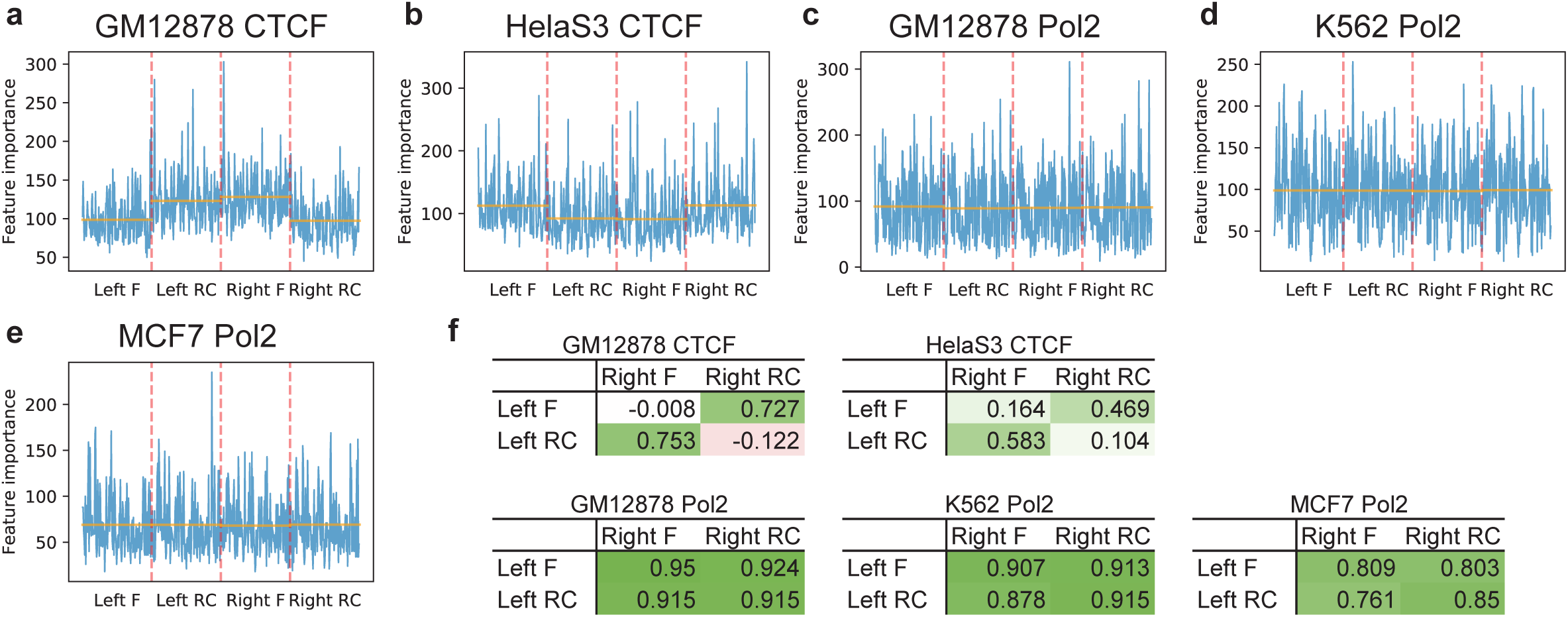
Sequence feature importance scores of gradient boosted trees trained on extended datasets. **a-e**, The importance scores of sequence features extracted from both directions (F: forward; RC: reverse complement) of the two anchors (left and right) by models trained on different datasets. The orange horizontal lines indicate average importance scores of the features from the strand of the anchor. **f**, Pearson correlations between feature importance scores of the two anchors.

**Figure 4:**
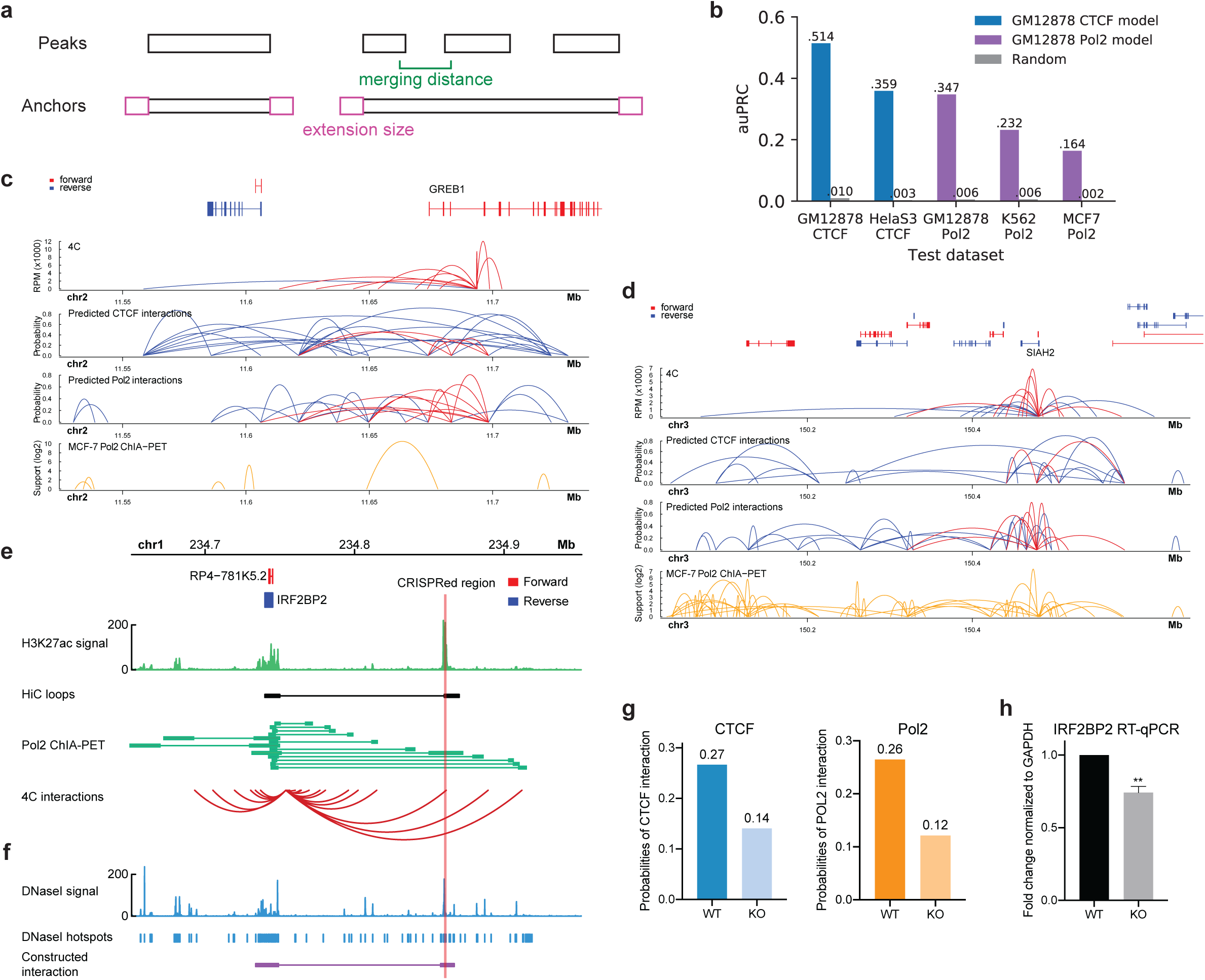
Performances of the from-DNase models and validations. **a**, Illustration of the two parameters, merging distance and extension size, used in constructing putative chromatin interactions anchors from open chromatin regions. **b**, Area under precision-recall curves of the from-DNase models. **c-d**, Validations of the predicted chromatin interactions by 4C-seq at *GREB1* and *SIAH2* gene regions in *MCF-7* cells. **e**, H3K27ac signal and chromatin interactions identified by Hi-C, Pol2 ChIA-PET and 4C-seq at the *IRF2BP2* promoter region in K562 cells. The CRISPR deleted enhancer region is highlighted by the vertical red line. **f**, DNase-seq signal, hotspots and the chromatin interaction constructed from the hotspots linking *IRF2BP2* and the target enhancer region. **g**, The predicted probabilities of CTCF- and Pol2-associated chromatin interactions and **h**, *IRF2BP2* expression before (WT) and after (KO) the deletion of the enhancer region by CRISPR.

For each model, we obtained and matched the position-weight matrices for all kernels on the first convolutional layer to known transcription factor binding motifs (Supplementary Figure 3). As expected, CTCF motif was captured by both CTCF models (Supplementary Figure 3a-b). Other than the CTCF motif, the remaining known transcription factor binding motifs learned by the two models were different. The patterns learned by Pol2 models showed more diversity and no matching transcription factor binding motif was shared among the three models (Supplementary Figure 3c-e). Interestingly, some of the transcription factors identified, such as ZNF143 in K562 and GATA3 in MCF-7, play important roles in the relevant cancer types^27, 28^.

### Convergent CTCF motifs are important for prediction of CTCF-associated open chromatin interactions

After extracting the sequence features from both the forward and reverse-complement sequences of the anchors, they were fed into the classifier to obtain a probability score that indicated how likely the pair of anchors were involved in a chromatin interaction. We obtained the feature importance scores of the gradient boosted trees trained and validated using a set of extended datasets that includes more negative samples than the distance-matched datasets (Methods, Supplementary Figure 4a-d). Distance was the most important feature in all models. We focused on the sequence features that were important for the prediction. Interestingly, in CTCF models the important sequence features were on different strands of the two anchors (Figure 3a-b), while Pol2 models did not show such pattern (Figure 3c-e). For the CTCF models, importance scores of features on different strands of the two anchors showed good correlation, while importance scores of features on the same strand of the two anchors did not show much correlation (Figure 3f). In contrast, the importance scores of features of Pol2 models were generally highly correlated regardless of the strand. The kernels on the last convolutional layer that generated the most important features in the extended CTCF models captured the CTCF motif (Supplementary Figure 4e-f), suggesting that convergent CTCF motifs were important for the prediction of CTCF-associated chromatin interactions. However, using only CTCF motif information for the prediction of CTCF-associated open chromatin interactions could not recapitulate the performance achieved by the convolutional neural network (Supplementary Figure 4g).

### Predicting chromatin interactions from open chromatin regions

The above models were trained and evaluated on known chromatin interactions. Without knowledge of chromatin interactions, as is the case for many clinical samples and cell types, the locations of the anchors would not be known. To be able to predict chromatin interactions between open chromatin regions, the models need to be able to predict chromatin interactions between anchors constructed from open chromatin regions. We tested different combinations of merging distances and extension sizes (Figure 4a) based on validation datasets and determined that the merging distance of 3000 bp and extension size of 1000 bp for the construction of anchors in GM12878 cells (Supplementary Figure 5a).

**Figure 5:**
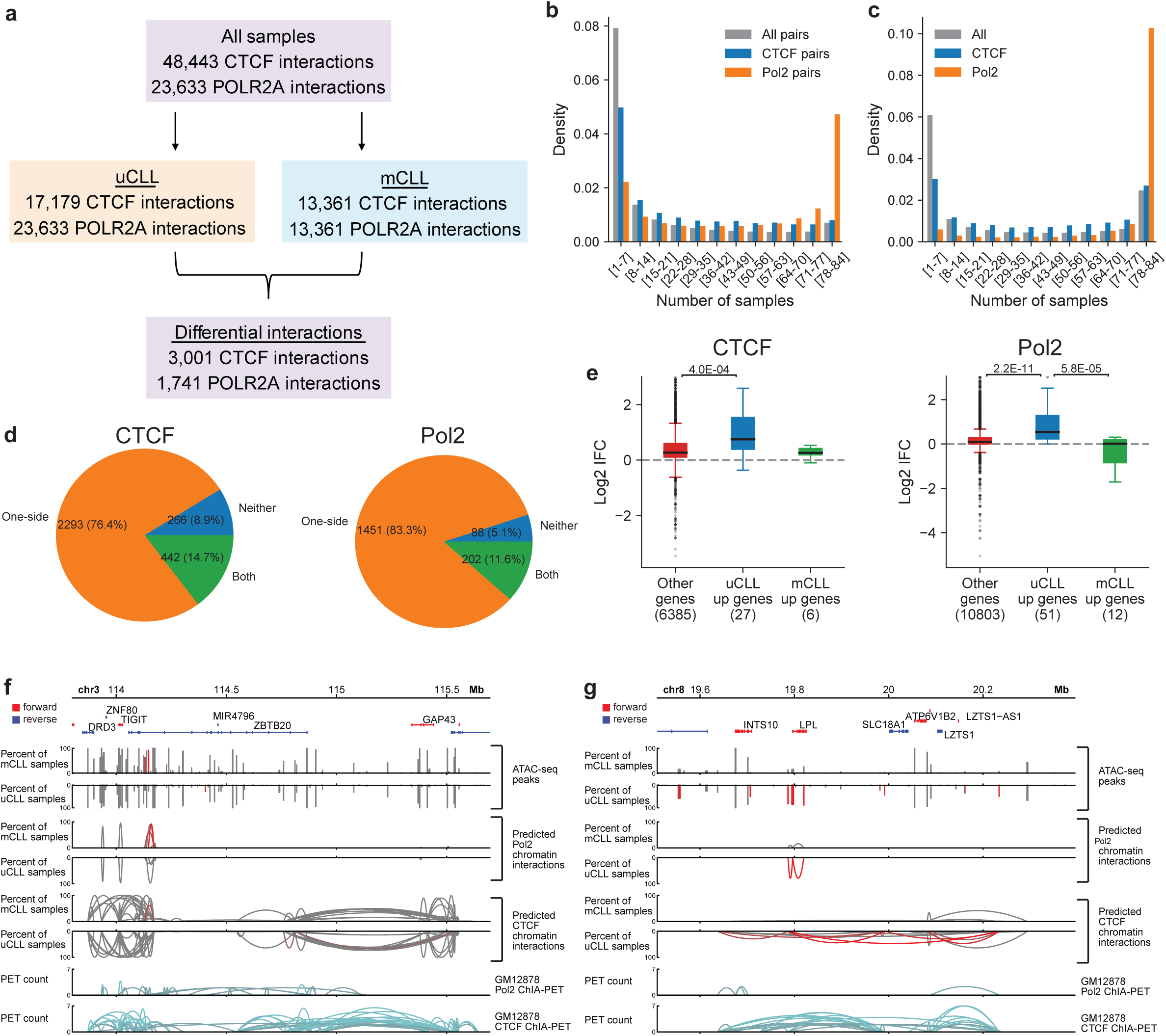
Predicted chromatin interactions in CLL samples. **a**, Summary of the predicted chromatin interactions in the 84 CLL samples and the differential chromatin interactions between uCLL and mCLL samples. **b**, Conservation analysis of predicted chromatin interactions in the CLL samples. All pairs: all possible pairs used for prediction. **c**, Uniqueness analysis of open chromatin regions that overlap with CTCF or Pol2 peaks from GM12878 cells in the CLL samples. All: all open chromatin regions. **d**, Distribution of differential CTCF and Pol2 chromatin interactions based on whether both anchors (Both), one anchor (One-side), or neither anchors (Neither) showed the same level of differences between uCLL and Mcll samples as the associated chromatin interaction. **e**, Association of differences in chromatin interactions between uCLL and mCLL samples with differentially expressed genes identified from a set of microarray samples. IFC: the fold change of the average number of chromatin interactions observed at the gene promoter in uCLL samples over that in mCLL samples. p-values were calculated using the Kruskal-Wallis test. **f-g**, Examples of genes, *ZBTB20* and *LPL*, whose different connectivity are associated with differences in distal regions. The red bars and curves indicate significantly different open chromatin regions and chromatin interactions based on Fisher’s Exact test.

The pairs generated between anchors constructed from open chromatin regions in GM12878 were used to train gradient boosted trees for both CTCF and Po l2 models (see Methods). The positive-to-negative ratios were about 1:122 for CTCF chromatin interaction labeled samples and 1:186 for Pol2 chromatin interaction labeled samples. The CTCF model achieved within-sample auPRC of 0.514 and the Pol2 model achieved auPRC of 0.347 (Figure 4b). In cross-sample evaluation, the CTCF model achieved auPRC of 0.359 on HelaS3 CTCF dataset and the Pol2 model achieved auPRCs of 0.232 and 0.164 on K562 and MCF-7 Pol2 datasets, respectively (Figure 4b). We were able to validate some of the predicted chromatin interactions in MCF-7 cells using 4C-seq (Figure 4c-d, Supplementary Figure 4b). Some of the validated chromatin interactions were not captured by the MCF-7 Pol2 ChIA-PET dataset.

We tested whether the models were able to pick up the genetic deletion of an enhancer region of *IRF2BP2* in K562 cells by CRISPR (Figure 4e-f). Interestingly, when the deleted sequences were removed from the input to the model, the predicted probabilities decreased for both CTCF- and Pol2-associated chromatin interactions between the enhancer and *IRF2BP2* (Figure 4g), which was consistent with the observation that deletion of the enhancer region led to decrease in the expression of *IRF2BP2* in K562 cells (Figure 4h).

### Exploring chromatin interactions in CLL samples

We applied the above models to 84 chronic lymphocytic leukemia (CLL) samples whose open chromatin profiles were available by ATAC-seq^24^. The CLL samples could be divided into two subtypes based on IGHV mutation status: 34 IGHV-unmutated CLL (uCLL) samples and 50 IGHV-mutated CLL (mCLL) samples. IGHV mutation status is an important prognostic biomarker in CLL, with mCLL being less aggressive^24^. A total of 48,443 CTCF-associated open chromatin interactions and 23,633 Pol2-associated open chromatin interactions were predicted based on the pooled open chromatin regions of all samples (Figure 5a). Pol2-associated chromatin interactions were better conserved across the CLL samples than CTCF-associated chromatin interactions (Figure 5b), which could be attributed to that open chromatin regions in the CLL samples that overlapped with GM12878 Pol2 peaks were better conserved than those overlapping with GM12878 CTCF peaks (Figure 5c). Using this set of ATAC-seq data in CLL samples, it was reported that regions with higher open chromatin signals in uCLL samples showed strong enrichment of binding sites of CTCF, RAD21 and SMC3^24^, which could also contribute to the high variability of CTCF chromatin interactions.

Variations in occurrences of chromatin interactions between the two subtypes of CLL were highly associated with variations in occurrences of anchor regions. Thus, using the predicted chromatin interactions, it was possible to separate mCLL and uCLL samples (Supplementary Figure 5a). Most differential chromatin interactions were associated with changes in the occurrence of one anchor (Figure 5d). There was a small portion of differential chromatin interactions whose anchors did not have the same level of changes as the chromatin interactions themselves between the two subtypes. In this set of differential chromatin interactions, the rate of co-occurrences of the two anchors within the same sample could change, contributing to the levels of changes in chromatin interactions (Supplementary Figure 5b).

Genes with higher expression in uCLL showed higher connectivity at the transcription start sites (Figure 5e, Supplementary Figure 5c). The differences in connectivity at transcription start sites were mostly associated with differences in the occurrences of the open chromatin regions at the transcription start sites (Supplementary Figure 5d). Nonetheless, there exist some genes, such as *WSB2* (Supplementary Figure 5e) and *ZBTB20* (Figure 5f), whose differences in connectivity were mostly associated with differences of the distal interacting regions. In addition, differences in connectivity at important CLL prognostic markers, such as *LPL* (Figure 5g), *ZAP70* (Supplementary Figure 5f), *ZNF667* (Supplementary Figure 5g), and *CD38* (Supplementary Figure 5h)^29-32^, were associated with differences in open chromatin regions at both transcription start sites and distal interacting regions.

## Discussion

We described a convolutional neural network, CHINN, that can extract sequence features and be coupled to classifiers to predict chromatin interactions between open chromatin regions using DNA sequences and distance. This approach only requires the use of open chromatin data and showed good generalizability on the same type of chromatin interactions across different cell types. Thus, it has the potential to be applied to large sets of clinical samples with limited biological materials. In addition, CHINN can discover sequence features that are important for predicting chromatin interactions, including shared features such as the CTCF motif and cell-type specific features such as GATA3 binding motif in MCF-7 and ZIC3 motif in HelaS3.

We showed that at resolutions limited by the experimental techniques, chromatin interactions between open chromatin regions could be predicted from 1-dimensional functional genomics data with reasonable accuracy. In distance-controlled experiments, our prediction method using functional genomics data performed better on Pol2 chromatin interactions but worse on CTCF chromatin interactions compared to sequence-based CHINN. Such differences could be attributed to the lower functional genomic complexity at CTCF binding sites and functional genomic data might fail to capture the convergent CTCF binding motifs often observed at CTCF-mediated chromatin interactions. On the other hand, Pol2 binding sites do not have distinctive DNA motifs, making it harder to predict Pol2 binding sites^22, 23^ and consequently harder to predict Pol2-associated chromatin interactions from DNA sequences. However, Pol2 binding sites are usually occupied by many other transcription factors, making it easier to predict Pol2-associated chromatin interactions using functional genomic data.

The application of CHINN models with gradient boosted tree classifiers to a cohort of CLL ATAC-seq samples revealed that there existed systematic differences in chromatin interactions involving important CLL prognostic genes, such as *LPL* and *CD38,* between the IGHV-mutated and IGHV-unmutated subtypes. These results suggest that differences in chromatin interaction landscapes between CLL subtypes could have important functional implications in CLL biology. In conclusion, the CHINN method may be useful in the future in understanding chromatin interactions in large cohorts of clinical samples and identifying chromatin interaction-based biomarkers.

## Data availability

4C-seq data are deposited in Gene Expression Omnibus (GSE135052).

## Code availability

Code and training data will be made available upon request.

## References

1. Babu, D. & Fullwood, M.J. 3D genome organization in health and disease: emerging opportunities in cancer translational medicine. Nucleus 6, 382–393 (2015).

2. Schottenfeld, D. in Gastrointestinal Oncology: Principles and Practice. (eds. D.P. Kelsen, J.M. JDaly, B. Levin, S.E. Kern & J.E. Tepper) (Lippincott Williams and Wilkins, Philadelphia; 2002).

3. Li, G. et al. Extensive promoter-centered chromatin interactions provide a topological basis for transcription regulation. Cell 148, 84–98 (2012).

4. Jin, F. et al. A high-resolution map of the three-dimensional chromatin interactome in human cells. Nature 503, 290–294 (2013).

5. Ma, W. et al. Fine-scale chromatin interaction maps reveal the cisregulatory landscape of human lincRNA genes. Nat Methods 12, 71–78 (2015).

6. Dowen, Jill M. et al. Control of Cell Identity Genes Occurs in Insulated Neighborhoods in Mammalian Chromosomes. Cell 159, 374–387 (2014).

7. Lieberman-Aiden, E. et al. Comprehensive mapping of long-range interactions reveals folding principles of the human genome. Science 326, 289–293 (2009).

8. Fullwood, M.J. et al. An oestrogen-receptor-alpha-bound human chromatin interactome. Nature 462, 58–64 (2009).

9. Dixon, J.R. et al. Topological domains in mammalian genomes identified by analysis of chromatin interactions. Nature 485, 376–380 (2012).

10. Lupianez, D.G. et al. Disruptions of topological chromatin domains cause pathogenic rewiring of gene-enhancer interactions. Cell 161, 1012–1025 (2015).

11. Guo, Y. et al. CRISPR Inversion of CTCF Sites Alters Genome Topology and Enhancer/Promoter Function. Cell 162, 900–910 (2015).

12. Rao, S.S. et al. A 3D map of the human genome at kilobase resolution reveals principles of chromatin looping. Cell 159, 1665–1680 (2014).

13. Cao, Q. et al. Reconstruction of enhancer-target networks in 935 samples of human primary cells, tissues and cell lines. Nature genetics 49, 1428–1436 (2017).

14. He, B., Chen, C., Teng, L. & Tan, K. Global view of enhancer-promoter interactome in human cells. Proceedings of the National Academy of Sciences of the United States of America 111, E2191–2199 (2014).

15. Roy, S. et al. A predictive modeling approach for cell line-specific long-range regulatory interactions. Nucleic acids research 43, 8694–8712 (2015).

16. Singh, S., Yang, Y., Poczos, B. & Ma, J. Predicting Enhancer-Promoter Interaction from Genomic Sequence with Deep Neural Networks. bioRxiv, 85241 (2016).

17. Whalen, S., Truty, R.M. & Pollard, K.S. Enhancer-promoter interactions are encoded by complex genomic signatures on looping chromatin. Nature genetics 48, 488–496 (2016).

18. Yang, Y., Zhang, R., Singh, S. & Ma, J. Exploiting sequence-based features for predicting enhancer–promoter interactions. Bioinformatics 33, i252–i260 (2017).

19. Zhu, Y. et al. Constructing 3D interaction maps from 1D epigenomes. Nat Commun 7, 10812 (2016).

20. Xi, W. & Beer, M.A. Local epigenomic state cannot discriminate interacting and non-interacting enhancer-promoter pairs with high accuracy. PLoS Comput Biol 14, e1006625 (2018).

21. Cao, F. & Fullwood, M.J. Inflated performance measures in enhancer–promoter interaction-prediction methods. Nature genetics (2019).

22. Zhou, J. & Troyanskaya, O.G. Predicting effects of noncoding variants with deep learning-based sequence model. Nat Methods 12, 931–934 (2015).

23. Alipanahi, B., Delong, A., Weirauch, M.T. & Frey, B.J. Predicting the sequence specificities of DNA-and RNA-binding proteins by deep learning. Nat Biotechnol 33, 831–838 (2015).

24. Rendeiro, A.F. et al. Chromatin accessibility maps of chronic lymphocytic leukaemia identify subtype-specific epigenome signatures and transcription regulatory networks. Nat Commun 7, 11938 (2016).

25. Consortium, E.P. An integrated encyclopedia of DNA elements in the human genome. Nature 489, 57–74 (2012).

26. Chen, T. & Guestrin, C. in Proceedings of the 22nd ACM SIGKDD International Conference on Knowledge Discovery and Data Mining 785-794 (ACM, San Francisco, California, USA; 2016).

27. Gonzalez, D. et al. ZNF143 protein is an important regulator of the myeloid transcription factor C/EBPalpha. The Journal of biological chemistry 292, 18924–18936 (2017).

28. Cimino-Mathews, A. et al. GATA3 expression in breast carcinoma: utility in triple-negative, sarcomatoid, and metastatic carcinomas. Hum Pathol 44, 1341–1349 (2013).

29. Kaderi, M.A. et al. LPL is the strongest prognostic factor in a comparative analysis of RNA-based markers in early chronic lymphocytic leukemia. Haematologica 96, 1153–1160 (2011).

30. Morabito, F. et al. Surrogate molecular markers for IGHV mutational status in chronic lymphocytic leukemia for predicting time to first treatment. Leuk Res 39, 840–845 (2015).

31. Rozovski, U. et al. Aberrant LPL Expression, Driven by STAT3, Mediates Free Fatty Acid Metabolism in CLL Cells. Mol Cancer Res 13, 944–953 (2015).

32. Crespo, M. et al. ZAP-70 expression as a surrogate for immunoglobulin-variable-region mutations in chronic lymphocytic leukemia. The New England journal of medicine 348, 1764–1775 (2003).

