## Supplementary Figures for "Predicting chromatin interactions between open chromatin regions from DNA sequences"

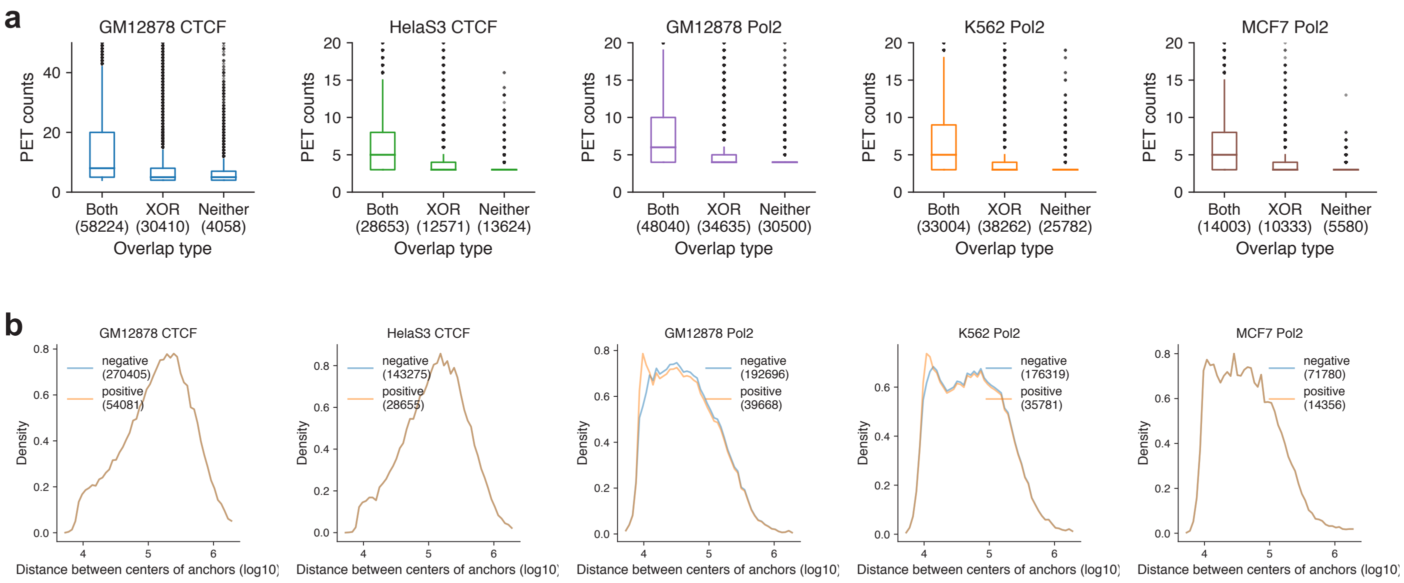


**Supplementary Figure 1: Characteristics of positive and negative samples in distance-matched datasets. a,** Comparison of the support of chromatin interactions based on whether both anchors (Both), only one anchor (XOR), or neither anchors (Neither) overlap with open chromatin regions. The number in brackets indicates the number of chromatin interactions in the category. **b,** Distance distributions of positive and negative samples in distance-matched datasets.

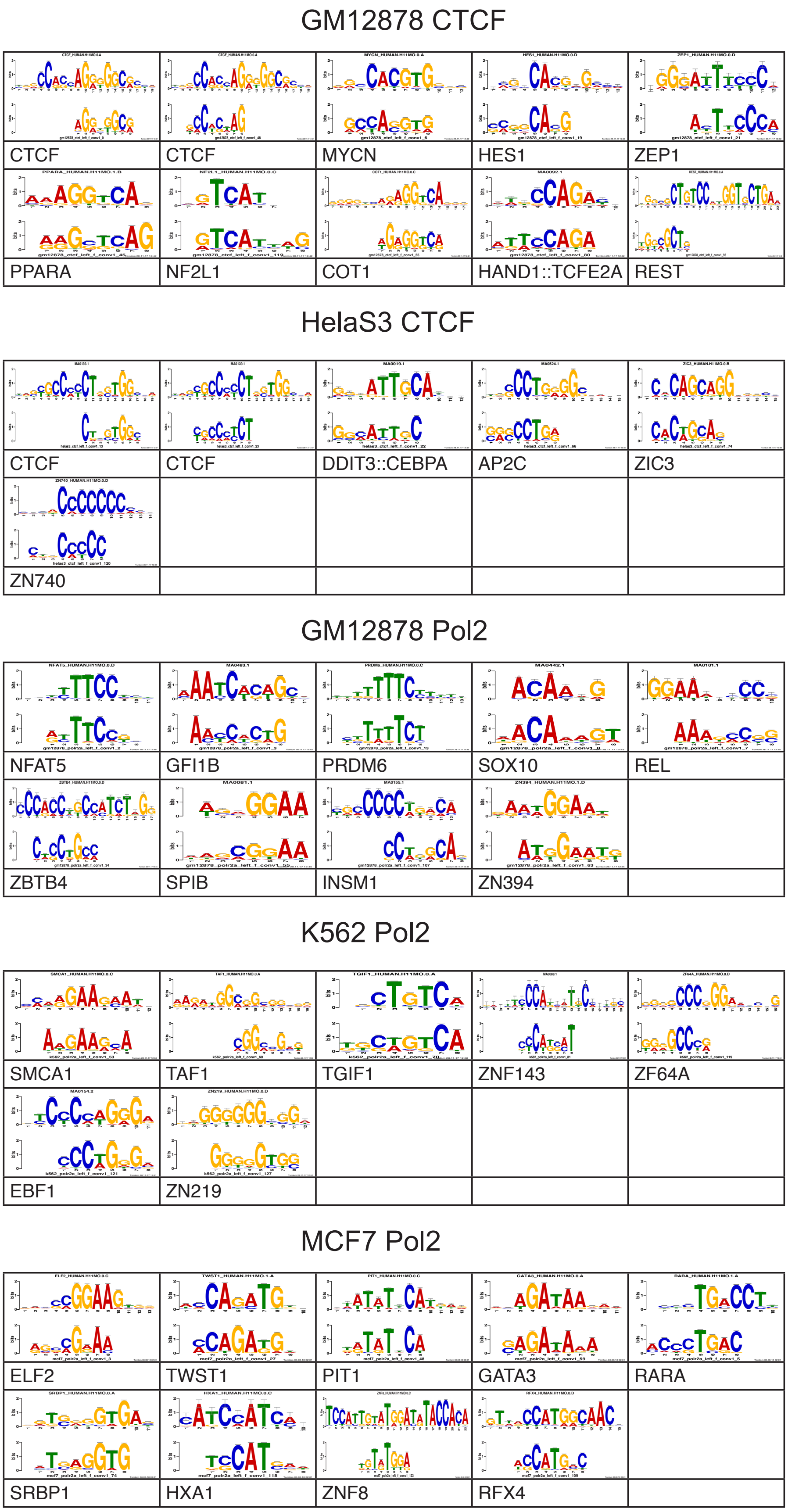


**Supplementary Figure 2: Convolutional layer 1 kernels of the five models that resemble known transcription factor binding motifs.**

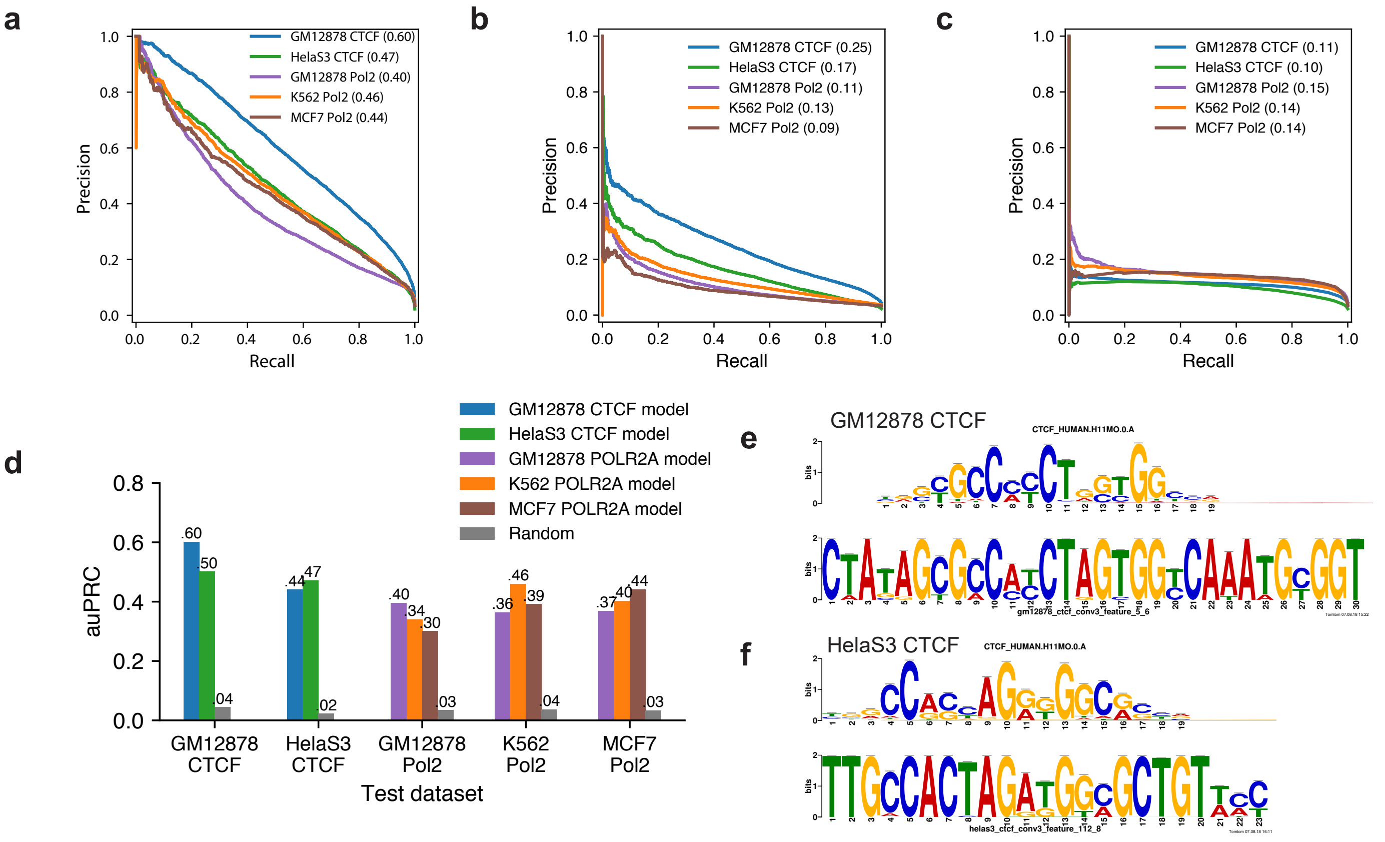


**g**

| auPRC | | Tested on GM12878 | Tested on HelaS3 |
|---|---|---|---|
| Trained on | GM12878 | 0.272 | 0.223 |
| | HelaS3 | 0.23 | 0.25 |

**Supplementary Figure 3: Sequence models on extended datasets. a-c,** Precision-recall curves of sequence models on extended datasets using (**a**) sequence features and distance; (**b**) only sequence features; and (**c**) only distance. d, Across-sample performancesof the sequence models on extended datasets. **e-f,** Top layer 3 kernels of CTCF models captured the CTCF motif. **g,** Performances of using only CTCF motif occurrences, motif orientations, and distance for prediction.

**a**

| CTCF auROC | Extension size (bp) 500 | 1000 | 1500 | 2000 | 2500 | 3000 |
|---|---|---|---|---|---|---|
| Merging size (bp) 500 | 0.933 | 0.942 | 0.948 | 0.952 | 0.953 | 0.952 |
| 1000 | 0.944 | 0.946 | 0.95 | 0.954 | 0.954 | 0.954 |
| 1500 | 0.95 | 0.952 | 0.951 | 0.953 | 0.953 | 0.953 |
| 2000 | 0.953 | 0.955 | 0.954 | 0.954 | 0.953 | 0.953 |
| 2500 | 0.954 | 0.956 | 0.956 | 0.954 | 0.953 | 0.953 |
| 3000 | 0.955 | 0.957 | 0.956 | 0.955 | 0.954 | 0.952 |

| Pol2 auROC | Extension size (bp) 500 | 1000 | 1500 | 2000 | 2500 | 3000 |
|---|---|---|---|---|---|---|
| Merging size (bp) 500 | 0.926 | 0.932 | 0.934 | 0.934 | 0.934 | 0.934 |
| 1000 | 0.937 | 0.94 | 0.94 | 0.939 | 0.939 | 0.938 |
| 1500 | 0.943 | 0.946 | 0.946 | 0.945 | 0.945 | 0.944 |
| 2000 | 0.947 | 0.948 | 0.948 | 0.947 | 0.947 | 0.946 |
| 2500 | 0.953 | 0.954 | 0.953 | 0.953 | 0.952 | 0.951 |
| 3000 | 0.954 | 0.954 | 0.953 | 0.953 | 0.952 | 0.951 |

**b**

**c**

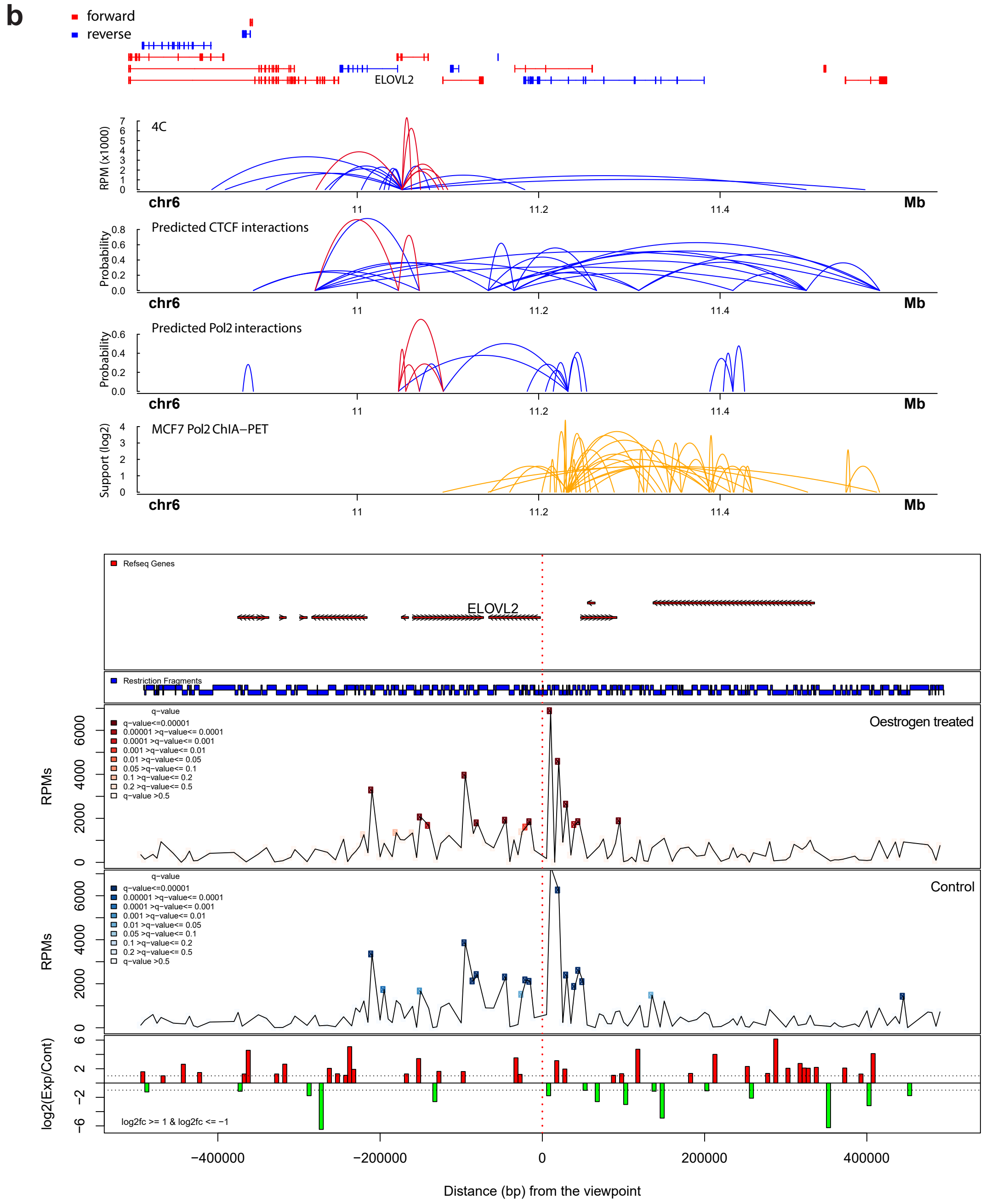


**Supplementary Figure 4a-c: Selection of parameters for from-dnase models and validations by 4C-seq in MCF-7 cells. a,** auROC of models trained on extended datasets based on different combinations of merging distances and extension sizes. **b,** Validations of predicted chromatin interactions by 4C-seq at *EVOVL2* gene locus. **c,** Comparison of 4C-seq chromatin interactions between oestrogen-treated and control MCF-7 cells at *ELOVL2* gene.

d

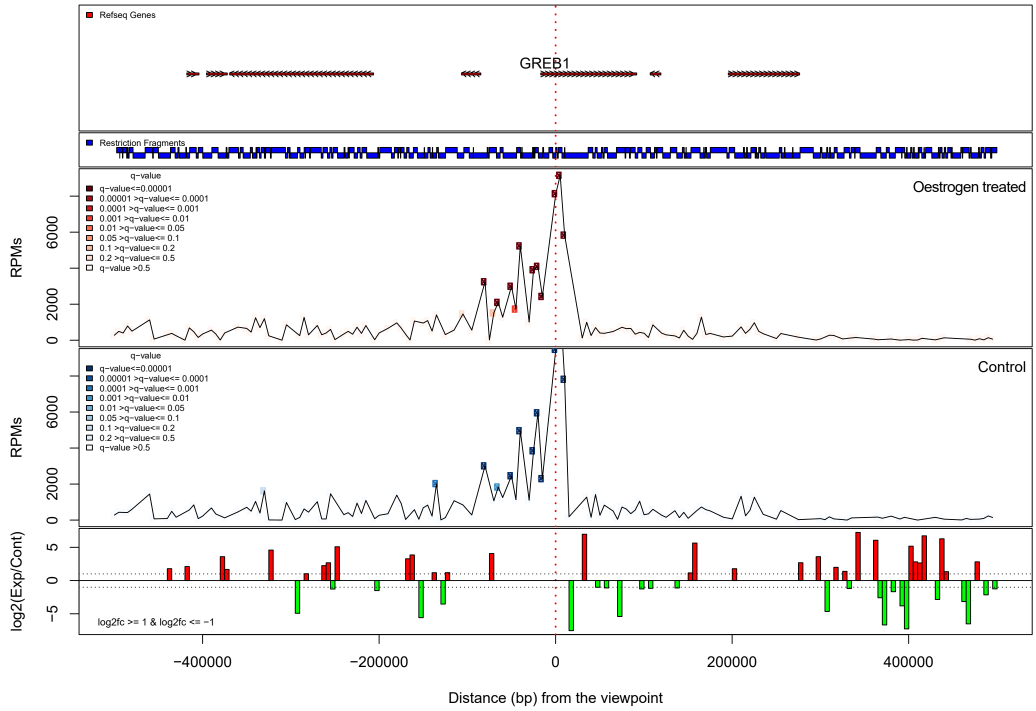


e

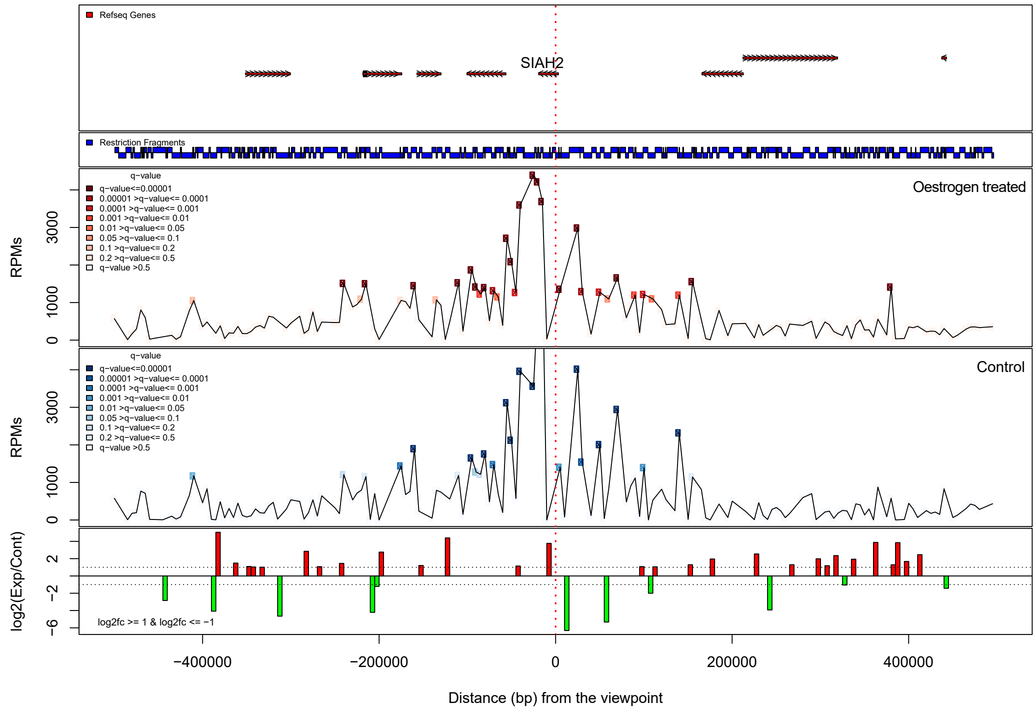


**Supplementary Figure 4d-e:** Comparison of 4C-seq chromatin interactions between oestrogen-treated and control MCF-7 cells at *GREB1* and *SIAH2* gene.

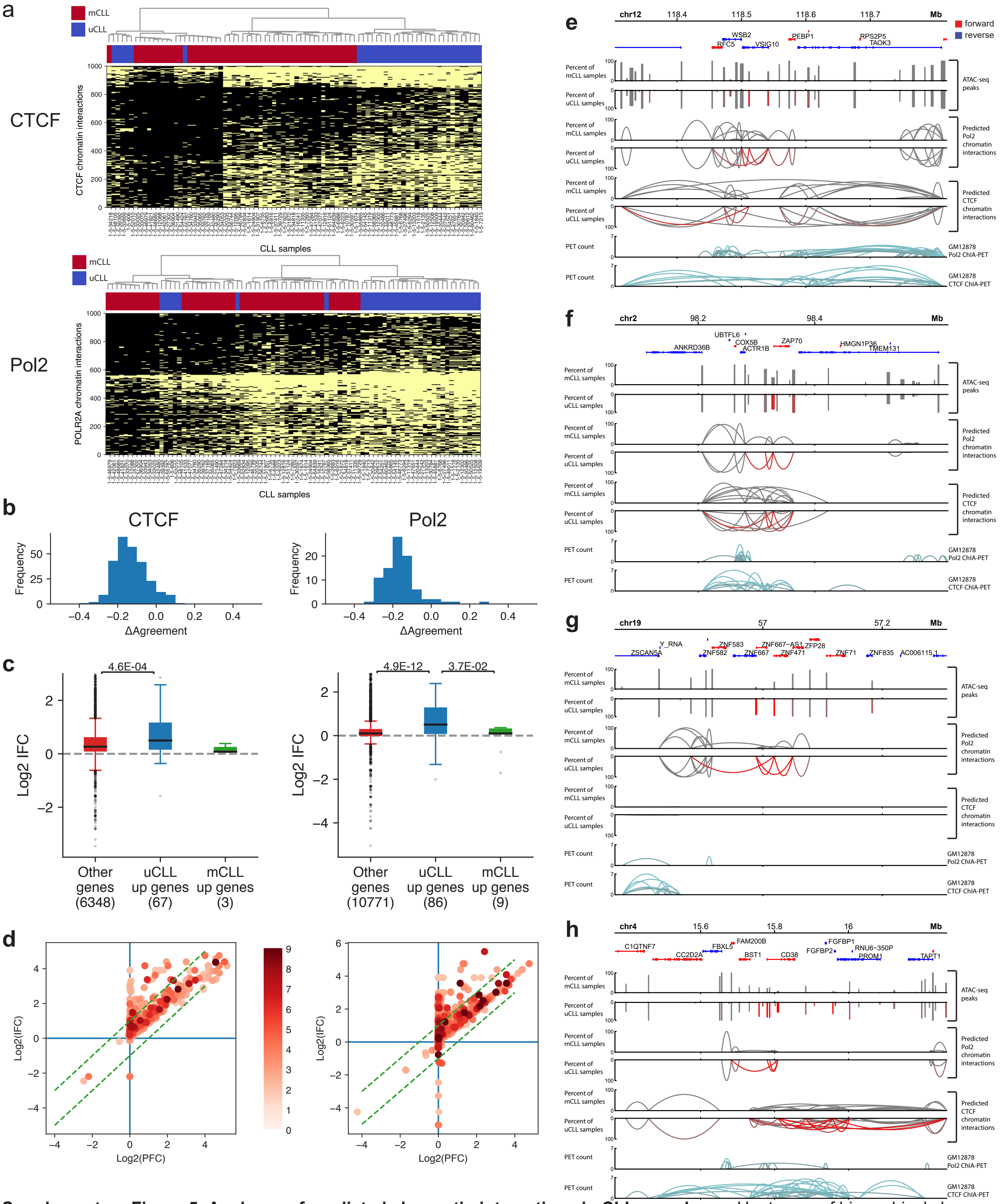


**Supplementary Figure 5: Analyses of predicted chromatin interactions in CLL samples. a,** Heatmaps of hierarchical clustering of CLL samples based on the 1000 most important chromatin interactions of the CTCF and Pol2 random forest classiers, respectively. Yellow indicates the presence of a chromatin interaction and black indicates the absence of a chromatin interaction. Red color represents mCLL samples and blue represents uCLL samples. **b,** Differences in agreement between anchors of differential CTCF and Pol2 chromatin interactions in the ‘Neither’ category between mCLL and uCLL samples. **c,** Association of differences in chromatin interactions between uCLL and mCLL samples with differentially expressed genes identified using RNA-seq data. IFC: the fold change of the average number of chromatin interactions observed at the gene promoter in uCLL samples over that in mCLL samples. p-values were calculated using Kruskal-Wallis test. **d,** The association between differences in chromatin interactions and differences in open chromatin peaks at gene promoter between uCLL and mCLL samples. PFC: the fold change of the proportions of samples the open chromatin peak at gene promoter is observed in uCLL samples over that in mCLL samples. The color bar indicates the -log(pvalue) of the most signicantly differential chromatin interaction at the promoter. **e-h,** Examples of genes whose different connectivity are associated with differences in distal regions. The red bars and curves indicate significantly different open chromatin regions and chromatin interactions based on Fisher’s Exact test.
